# Behavioural analysis of loss of function zebrafish supports *baz1b* as master regulator of domestication

**DOI:** 10.1101/2022.02.02.478842

**Authors:** Jose V. Torres-Pérez, Sofia Anagianni, Aleksandra M. Mech, William Havelange, Judit García-González, Scott E. Fraser, Giorgio Vallortigara, Caroline H Brennan

## Abstract

Domestication is associated with both morphological and behavioural phenotypic changes that differentiate domesticated species from their wild counterparts. Some of the traits are those purposely targeted by the selection process, whilst others co-occur as a result of selection. The combination of traits is referred to as the ‘domestication syndrome’ and their shared characteristics has given rise to the neural crest domestication syndrome (NCDS) hypothesis. According to this hypothesis, the phenotypic changes are a consequence of a selection towards animals with mild underdevelopment of the neural crest. A similar mechanism is suggested to affect some species of “self-domesticated” animals, including humans. Recent research supports a role for *BAZ1B*, one of the haplo-insufficient genes in Williams syndrome (WS), in the evolution of craniofacial features of modern humans. Interestingly, WS recapitulates some of the traits observed in the domestication syndrome, including hypersociability and reduced facial bones. However, the evidence linking *BAZ1B* to behavioural phenotypes associated with domestication is presumptive. Here, we use zebrafish as a model to test the hypothesis that *baz1b* loss-of-function leads to both morphological and behavioural phenotypes associated with the domestication syndrome by influencing the development of the neural crest. Our research provides further evidence supporting the NCDS hypothesis and *baz1b’s* role in this process.

## Introduction

Animal domestication represents an accelerated and directed artificial selection process towards, among others, increased tameness (1). Although the phenotypic changes that distinguish domesticated species from their wild ancestors/counterparts are taxon-specific, there is a set of recurrent morphological, behavioural and physiological traits within the different domesticated species. These features are described by the so-called “Domestication Syndrome” (DS) and include increased sociability, lower aggressiveness and reduced fear- and stress-related behaviours (1, 2). Additionally, DS encompasses other traits that do not seem directly selected but appear linked as byproducts of the above such as reduced craniofacial features, small facial bones and teeth, and depigmentation.

The existence of conserved DS-related traits supports the neural crest domestication syndrome (NCDS) hypothesis, which suggests that mild neurocristopathy (defects in neural crest-derived tissues) is the developmental feature driving the first stages of the process of domestication (1). According to this hypothesis, mild neural crest deficits result in less/slower input of cells to the target structures, which ultimately accounts for: reduced craniofacial features, hypofunction of adrenal and sympathetic ganglia, extended immaturity period of the Hypothalamus-Pituitary-Adrenal (HPA) axis, and a significant size reduction in certain components of the forebrain’s limbic system (3). Mild neurocristopathy is also associated with an extended immaturity period that supports the existence of a longer “socialization window” during early life (3–5). Therefore, the NCDS hypothesis offers a unitary explanation for all the DS-traits with a predicted genetic component involving the development of the neural crest. However, the NCDS hypothesis remains under debate as evidence demonstrating a causal link between underdevelopment of the neural crest and DS is scarce (6, 7) and most domesticated species only present a subset of DS-predicted features (4, 8).

Some species, including humans and other primates, seem to have also undergone a process of domestication without the presence of an external domesticating agent (self-domestication), which has favoured larger social groups and intraspecific interactions (9). Recent research supports the existence of a core mechanism for both domestication and self-domestication in favour of the NCDS hypothesis: In marmoset monkeys, a positive correlation has been found between the degree of vocal cooperation and the size of an individual’s white facial patch, thus establishing a link between behavioural (vocal co-operation) and morphological (change in pigmentation) DS-associated phenotypes (10). Paleogenomic analysis points to a role for the bromodomain transcription factor BAZ1B in the evolution towards reduced facial features, a key aspect of DS, in modern humans (11).

Baz1b is a chromatin remodeler that in both neural crest stem cells and *Xenopus* embryos regulates the balance between neural crest precursors and migratory cells (11, 12). Interestingly, *BAZ1B* is also one of the 28 genes affected in William’s syndrome (WS), an autosomal disease caused by haplo-insufficiency of a portion of chromosome 7, that recapitulates many aspects of the DS including facial features and pro-social behaviours (5, 13, 14). In contrast, duplication of the same chromosomic region results in autism-like disorders (15, 16). These findings, together with the paleogenomic data, raise the possibility that variations in *BAZ1B* underlie both neurological and behavioural changes associated with DS.

Here, we tested the hypothesis that deficits in the *baz1b* gene are causally related to both morphological and behavioural changes associated with domestication using zebrafish *(Danio rerio)*. Zebrafish have emerged as a novel organism in which to assess different aspects of behavioural neuroscience. The relative ease of generating genetic mutants, the conserved neuronal circuitry (17) and the fact that around 80% of genes associated with human diseases have a corresponding orthologue (18), make zebrafish an ideal model in which to study the genetics and neuronal circuitry underlying behaviour. Zebrafish can be of use to identify the effect of individual genes which are part of a genomic region with high linkage disequilibrium, such as *baz1b* in WS. Key to this study, zebrafish are shoaling species that have proved useful to assess gene expression changes associated with vertebrate domestication (19).

Our research shows that zebrafish *baz1b* loss of function (LoF) mutants show mild under-development of the neural crest at larval stages, and distinctive craniofacial features and reduced anxiety-associated phenotypes later in life. During the ontogeny of social behaviour, *baz1b* LoF mutants show an increased inclination to interact with conspecifics. Thus, *baz1b* LoF recapitulates both morphological and behavioural phenotypes associated with the DS in zebrafish supporting the NCDS hypothesis and confirming a key role of *baz1b* in this process. Identification of the prominent role of *baz1b* in domestication can help disentangle the genetic network driving this evolutionary process and, indirectly, pathways affecting anxiety and socially driven disorders.

## Results

### CRISPR/Cas9 induction of stable *baz1b* LoF zebrafish lines

We used the CRISPR/Cas9 system to induce mutations in the *baz1b* gene. Two lines were established, one, *baz1b^del44^*, carrying a 44-base pair (bp) deletion at exon 5, and the other, *baz1b^ins35^*,with a 35 bp insertion at exon 5. Both mutations result in the introduction of a premature stop codon (figure 1A, supplementary figure 2A). Quantification of *baz1b* mRNA levels by qPCR (figure 1B, supplementary figure 2B) revealed a significant difference in expression between genotypes in both lines *(baz1b^del44^*: F (2, 6) = 7.415, *p* = 0.0239; *baz1b^ins35^*: F (2, 6) = 21.97, *p* = 0.0017). Post-hoc analysis of the *baz1b^del44^* line showed that both heterozygous (HET, *p* = 0.0417) and homozygous (HOM, *p* = 0.0318) fish had significantly lower expression of *baz1b* than wild type (WT) siblings, while there was no significant difference between HET and HOM (*p* = 0.9725). On the *baz1b^ins35^* line, post-hoc analysis showed that HOM had significantly lower expression than WT (*p* = 0.0014) while HET failed to reach significance (vs. WT: *p* = 0.0638; vs. HOM: *p* = 0.0225).

**Figure 1.**
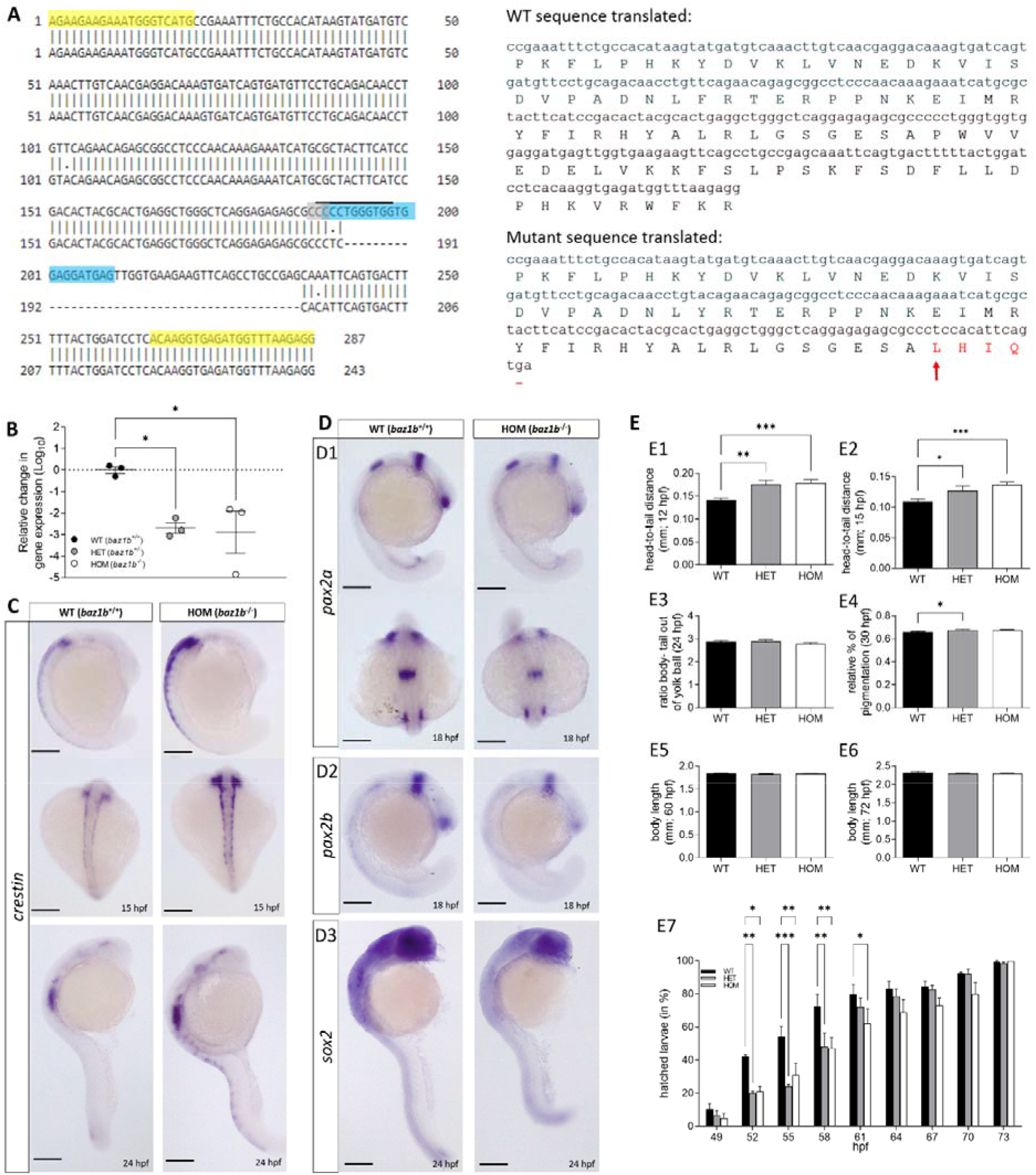
Deficits in *baz1b* lead to mild neurocristopathy in zebrafish. **A)** Left panel shows a DNA blast for the portion of *baz1b’s* exon 5 amplified by PCR used for genotyping between WT (above) and mutants (below, 44 bp deletion). Primers used are highlighted in yellow, crRNA site in blue, PAM site in grey and restriction enzyme over-lined. Right panel shows the in-frame translation to amino acids (aa) sequences in WT (above) and mutants (below) of the same exon 5 portion. Arrow indicates mutation starting site, changed aa are in red and stop codon marked as a dash (-). **B)** Relative change in gene expression (log_10_) assessed by qPCR between 5 dpf larvae from each genotype showing that in both HET and HOM the expression of *baz1b* is significantly diminished with respect to WT. Figure shows individual values (N = 3 per genotype, each representing a group of 16 larvae combined) and mean ± standard error mean (SEM). **C)** Whole larvae *in situ* hybridization (WISH) against *crestin* for WT and HOM at 18 and 24 hpf. **D)** WISH against *pax2a*, *pax2b* and *sox2*, genes regulated by *baz1b*. **E)** Developmental comparison between the three phenotypes for the head to tail distance at 12 hpf (E1) and at 15 hpf (E2), ratio body/tail-out-of-yolk-ball at 24 hpf (E3), relative percentage of pigmentation at 30 hpf (E4), body length at 60 hpf (E5) and at 72 hpf (E6), and percentage of hatched larvae between 49 to 73 hpf (E7). Graphs show mean ± SEM. In all cases: * *p* < 0.05; ** *p* < 0.01; *** *p* < 0.001. Scale bars represent 2 μm.

Since the primers target a region of the mRNA upstream from the mutation site, the decrease in functional mRNA suggests that the mRNA surveillance pathway interpreted the premature stop codon as aberrant and targeted the mRNA for degradation (20), consistent with establishment of a LoF line. As *baz1b^del44^* showed the most significant reduction in mRNA we have focused on analysis of this line. However, *baz1b^ins35^* showed similar morphological and behavioural differences as reported in supplementary figures (supplementary figures 2-5 and 7).

### Baz1b LoF leads to mild developmental delay at early stages

The expression of *baz1b* and neural crest induction begins during gastrulation (21), and the interaction between tissue types plays an important part in early development. Thus, we examined the impact of *baz1b* LoF on zebrafish development by assessing multiple morphological characteristics at different developmental points (figure 1E). At 12 hours post-fertilization (hpf), there was a significant difference in the head-to-tail distance between genotypes (H (2) = 16.45, *p* = 0.0003), with both HET (*p* = 0.0021) and HOM (*p* = 0.0008) embryos showing significantly larger distances than WT, suggesting delayed development. Similar differences were observed at 15 hpf (F (2, 78) = 7.480, *p* = 0.0011; HET vs. WT: *p* = 0.0466; HOM vs. WT: *p* = 0.0007).

No significant differences in overall body length were detected at 24 hpf or later stages of development (figure 1E). Nonetheless, a significant difference in the relative percentage of pigmentation was observed between genotypes at 30 hpf (F (2, 42) = 3.840, *p* = 0.0294) with post-hoc comparison describing a significant increase in HET (*p* = 0.0448) vs. WT. No other pairwise comparisons were significant. Additionally, analysis of hatching rate showed a significant interaction between time and genotype (F (16, 48) = 3.292, p = 0.0007). Post-hoc pairwise comparison revealed that the percentage of hatching was higher in WT than mutant siblings between 52 to 61 hpf (52 hpf: WT, vs. HOM: *p* = 0.0106, vs. HET: *p* = 0.0067; 55 hpf: WT, vs. HOM: *p* = 0.0037, vs. HET: *p* = 0.0002; 58 hpf: WT, vs. HOM: *p* = 0.0016, vs. HET: *p* = 0.0024; 61 hpf: WT, vs. HOM: *p* = 0.0345) but did not differ at later time points.

Changes with a similar directionality were observed in the *baz1b^ins35^* (supplementary figure 2).

### Baz1b LoF zebrafish show mild neurocristopathy

As Baz1b is implicated in neural crest development, we examined the development of the neural crest in whole larvae *in situ* hybridization (WISH) on larvae matched for developmental stage. The expression of *crestin*, a pan-neural crest marker of pre-migratory cells (22), was more widespread in the mutants than WT siblings at all developmental stages examined (figure 1C). The expression of genes involved in the differentiation and migration of neural crest, *pax2a, pax2b, sox2* (figure 1D) and *epha4* (supplementary figure 1), was reduced (12, 23–26). The reduced expression of *pax2a/b* and *epha4* was most evident in anterior areas, including the hindbrain and the optic vesicles. In contrast, the expression of *sox10*, a gene important for the formation and multipotency of the neural crest (27), and *foxd3*, implicated in neural crest maintenance (28), were unaffected (supplementary figure 1).

Zanella and colleagues (11) demonstrated that three genes, *ROBO1*, *ROBO2* and *FOXP2*, which are important for processes of vocal learning, a neurological feature also associated with domestication (10), are regulated by BAZ1B in humans. Therefore, we assessed how *baz1b* LoF affected the expression of these genes in zebrafish larvae. No visible differences in the pattern of expression of *robo1, robo2* and *foxp2* were observed between the different genotypes (supplementary figure 1).

The *baz1b^ins35^* line showed similar changes (supplementary figure 2).

### Baz1b LoF zebrafish have altered craniofacial features

One of the most recurring characteristics of DS is the size reduction of craniofacial features. In most species of mammals, domestication has led to shorter snouts, reduced teeth size and underdeveloped ear bones (1, 4). Paleogenomic data support a role for *BAZ1B* in this process in humans (11), and WS individuals, which lack a copy of *BAZ1B* among other genes, recapitulate DS-like facial features (13, 14). Therefore, we investigated whether *baz1b* LoF also induced craniofacial differences in zebrafish.

Larvae of 5 dpf from all genotypes were stained with alcian blue to measure their bone structures (figure 2A). Statistical analysis revealed that the ratio |width of the ceratohyal (CH) bone: length from Mechkel’s cartilage (M) to CH| was statistically different (H (3) = 12.53, *p* = 0.0019) and multiple pairwise post-hoc comparison revealed this to be driven by a decrease in HOM (*p* = 0.0015) when compared to WT siblings. Additionally, the ratio |length of M: length from M to CH| was also significantly different (H (3) = 14.87, *p* = 0.0006) with multiple pairwise post-hoc comparisons showing a reduction in both HET (*p* = 0.0388) and HOM (*p* = 0.0006) compared to WT. No other comparisons were significantly different.

**Figure 2.**
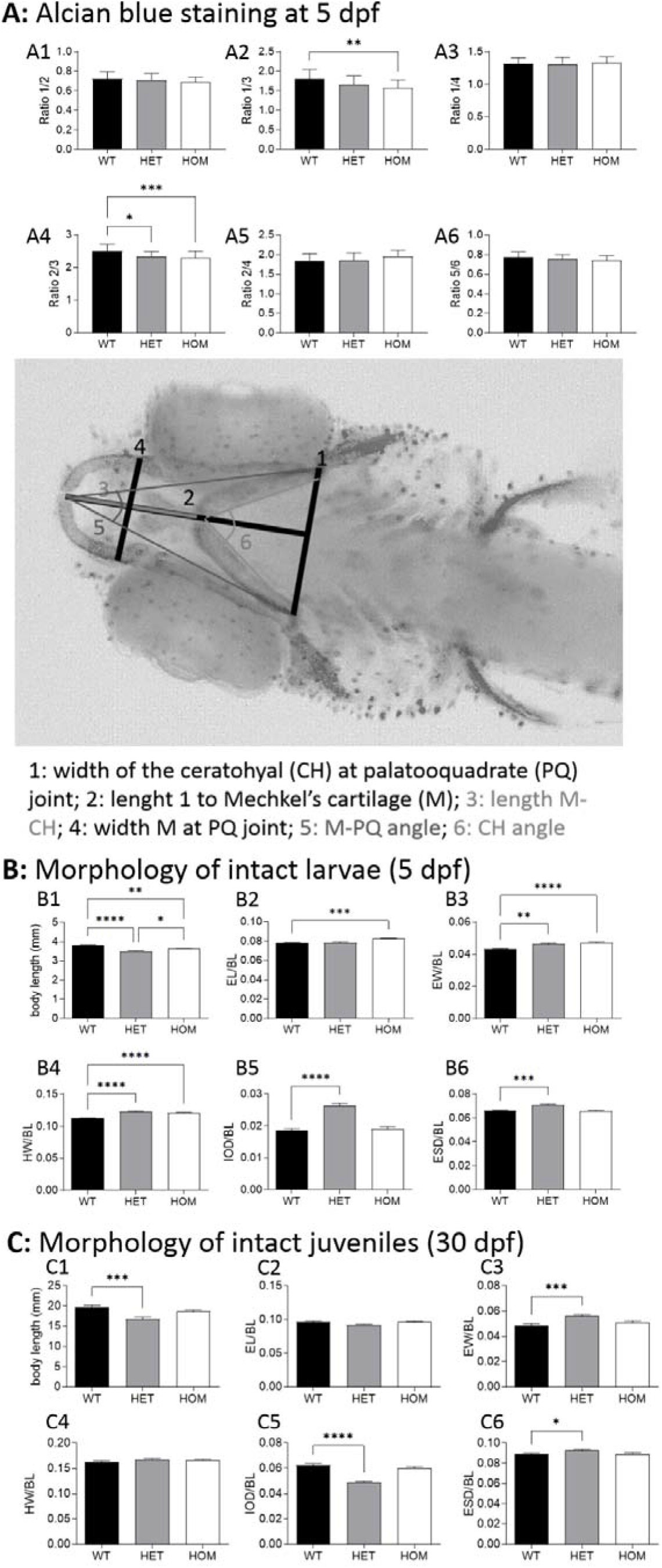
Baz1b LoF zebrafish have altered craniofacial features. **A)** Comparison between genotypes for the cranial features (outlined in the picture below) observed at 5 dpf by alcian blue staining: ratio 1/2 (A1), ratio 1/3 (A2), ratio 1/4 (A3), ratio 2/3 (A4), ratio 2/4 (A5) and ratio 5/6 (A6). **B)** Morphological comparisons of intact 5 dpf for the three genotypes: body length (BL, in mm; B1), eye length (EL) normalised to BL (B2), eye width (EW) normalised to BL (B3), head width (HW) normalised to BL (B4), inter-ocular distance (IOD) normalised to BL (B5) and eye-snout distance (ESD) normalised to BL (B6). **C)** Morphological comparison of intact juvenile fish (30 dpf). Graphs show mean ± SEM. In all cases: * *p* < 0.05; ** *p* < 0.01; *** *p* = 0.001; **** *p* < 0.001

To confirm that the changes in cranial bones were leading to facial variations (affecting the final shape of the soft tissues above the bones), we also performed morphological analysis of whole unstained larvae at 5 dpf (figure 2B). Firstly, we observed a significant difference in total body length (BL) between genotypes (H (3) = 39.12, *p* < 0.0001) with both HET and HOM showing reduced BL compared to WT siblings (HET: *p* < 0.0001, HOM: *p* = 0.0021). Reduction in BL was more pronounced in HET (HET vs. HOM: p =0.0130). This shortening in BL could be explained by a reduced input of neural crest cells which might have impacted the total length of the larvae, similarly to that reported in the *sox2*-morphants (29). Nonetheless, other researchers have pointed out that BL alone is not sufficient to assess differences due to the great range of size variability at any given age (30). Thus, the analysis of the additional partial body measurements (below) was done after standardising by BL.

We observed a significant change in eye length (EL, F (2, 57) = 9.465, *p* = 0.0003) and eye width (EW, F (2, 57) = 12.54, *p* < 0.0001) between genotypes, which post-hoc analysis showed to be driven by HOM having an increase for both (EL: *p* = 0.0009; EW: *p*< 0.0001) and HET only for EW (p = 0.0011) when compared to WT. Additionally, total head width (HW) was found to be significantly different between genotypes (F (2, 57) = 35.05, *p* < 0.0001) with both HET (*p* < 0.0001) and HOM (*p* < 0.0001) showing a significant increase when compared to WT. Furthermore, a significant difference was also observed regarding the inter-ocular distance (IOD, F (2, 57) = 40.35, *p*< 0.0001) and the eye-snout distance (ESD: F (2, 57) = 14.36, *p* < 0.0001), which in both cases post-hoc analysis revealed to be driven by a significant increase in the HET compared to WT (IOD: *p* <0.0001; ESD: *p* = 0.0001).

At 30 dpf, fish were also assessed on their maturation scores (30). These maturation scores rely on four external morphological characteristics (pigment pattern, tail fin, anal fin, and dorsal fin morphology) to determine their larval stage. No significant difference in the maturation scores was observed between the three genotypes (F (2, 57) = 1.013, *p* = 0.3695; mode in all genotypes was 4). In addition to assessing maturation, we analysed corresponding craniofacial measurements at 30 dpf, similarly as done at 5 dpf, to assess if changes were maintained at later stages (figure 2C). There was a significant difference in BL (F (2, 57) = 9.538, *p* = 0.0003) and post-hoc comparison revealed it to be driven by a reduction in HET (*p* = 0.0002) when compared to WT. Significant differences in EW were observed by genotype (F (2, 57) = 9.084, *p* = 0.0004) with HET showing significantly wider eyes than WT (*p* = 0.0004). As described at 5 dpf, there was a significant effect of genotype on IOD (F (2, 57) = 36.06, *p* < 0.0001) and ESD (F (2, 57) = 4.454, *p* = 0.0159). Pairwise post-hoc analysis showed those changes to be driven by HET. While ESD was increased in the HET when compared to WT (*p* = 0.0263), the IOD was diminished in the HET (*p* < 0.0001) compared to WT. No other significant differences were observed.

Data related to *baz1b^ins35^* LoF show differences with respect to the *baz1b^del44^* LoF (opposing directionality, supplementary figure 3), suggesting that the two lines could be useful for different types of study.

### Loss of *baz1b* diminishes stress response and habituation in larval zebrafish

As *baz1b* LoF affected maturation of the neural crest and facial morphology consistent with the NCDS hypothesis, we next examined whether *baz1b* LoF also showed behavioural differences associated with DS. Larvae from the three possible genotypes were assessed for stress reactivity at 5 dpf using a forced light-dark transition (FLDT) assay and response to brief flash of light, or acoustic startle (31).

Analysis of the FLDT assay revealed that *baz1b* LoF zebrafish habituate more quickly than WT siblings suggesting a diminished stress response (Figure 3). During the initial basal stage of the FLDT assay, before the alternating light/dark cycles there was a significant effect of genotype on locomotion (F (2) = 8.62, *p* = 0.013), where HET travelled shorter distances than the WT (*p* = 0.0126). Although HOM showed a similar reduction in overall distance travelled, they did not reach significance when compared to WT (*p* = 0.0793). The rate of increase in locomotion during the light periods, assessed as the combined slopes from the three light periods (first: minute 10 to 20, second: minute 30 to 40, third: minute 50 to 60) revealed a significant effect of time-genotype interaction (slope) (F (2) = 229.84, *p* < 0.001). Difference was observed between the mutant and WT, with the slope of HET and HOM being significantly steeper across the three light periods (HET: *p* = 0.033; HOM: *p* = 0.028). Dark periods, measured as a sum of slopes in dark periods that followed the light periods (first: minute 20 to 30, second: minute 40 to 50, third: minute 60 to 70) also revealed significant effect of time-genotype interaction (F (2) = 7.94, *p* =0.019) such that HET recovered significantly faster than WT (*p* = 0.024). There was no difference between HOM and WT (*p* = 0.189).

**Figure 3.**
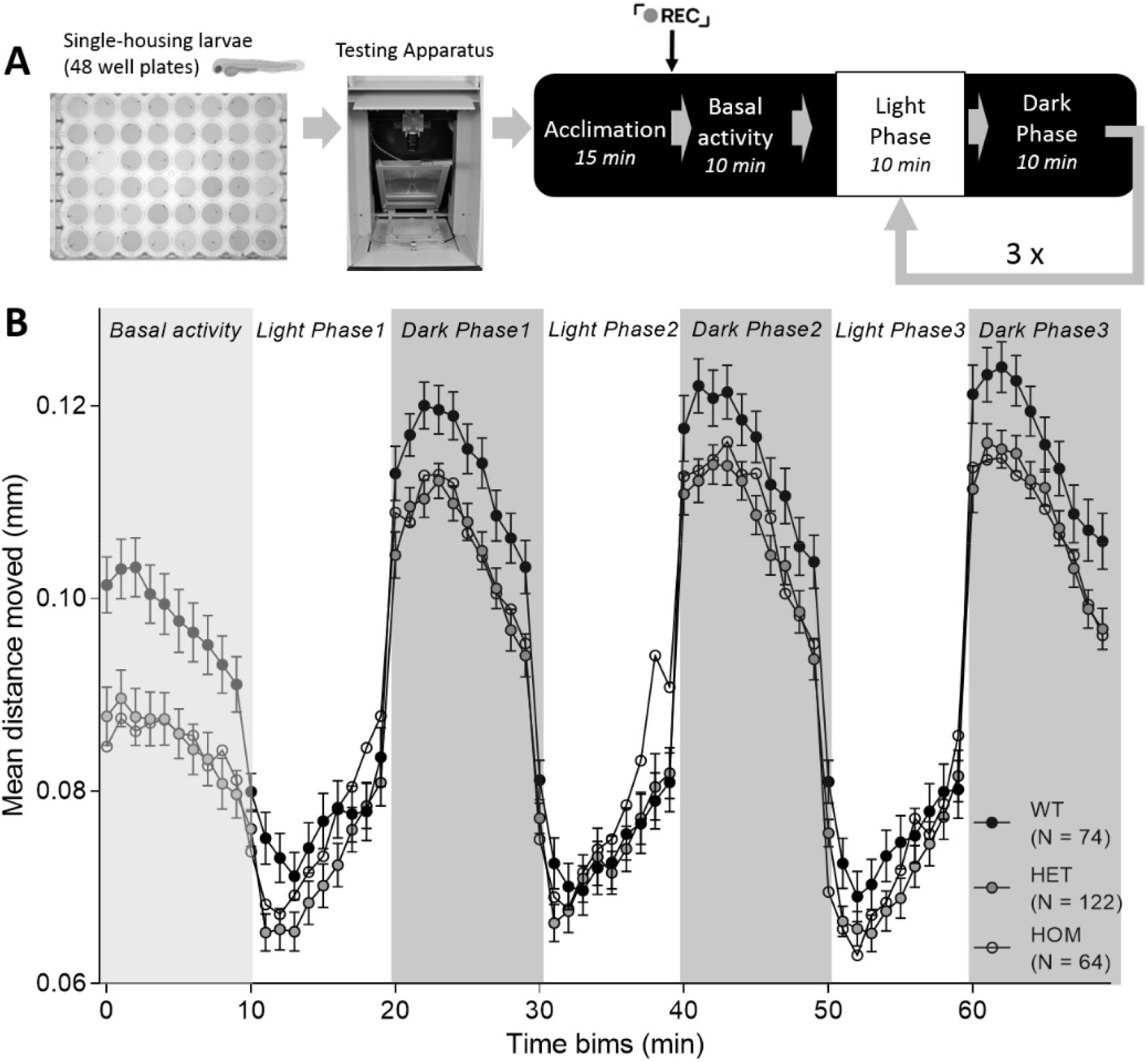
Loss of *baz1b* leads to faster habituation in the forced light dark transition (FLDT) assay. **A)** Diagram of the assay. **B)** Mean distance moved per genotype during the baseline and the three light-dark cycles (10 minutes each). Note that both HET and HOM describe a steeper slope than WT sibling during the light phases. Data shows mean ± SEM.

Similar to the FLDT assay, analysis of the flash of light paradigm showed that *baz1b* LoF also lead to a reduced stress reactivity in these assays (Figure 4B). The analysis of the baseline before the flash of light did not reveal significant differences between genotypes (*p* > 0.05). On transition from light to dark, in this assay, fish movement rapidly increases but the genotype wasn’t an indicator of the height of the jump (*p* > 0.05). However, there was a significant effect of time-genotype interaction on the slope of recovery following the flash of light (F (2)= 7.509, *p* = 0.023) between the HOM and WT (*p* = 0.0087). No other comparisons were significant. We next assessed locomotion after recovery from the flash of light and before the acoustic stimuli started. As in the first basal period, there was no significant difference between the genotypes in locomotion (*p* > 0.05).

**Figure 4.**
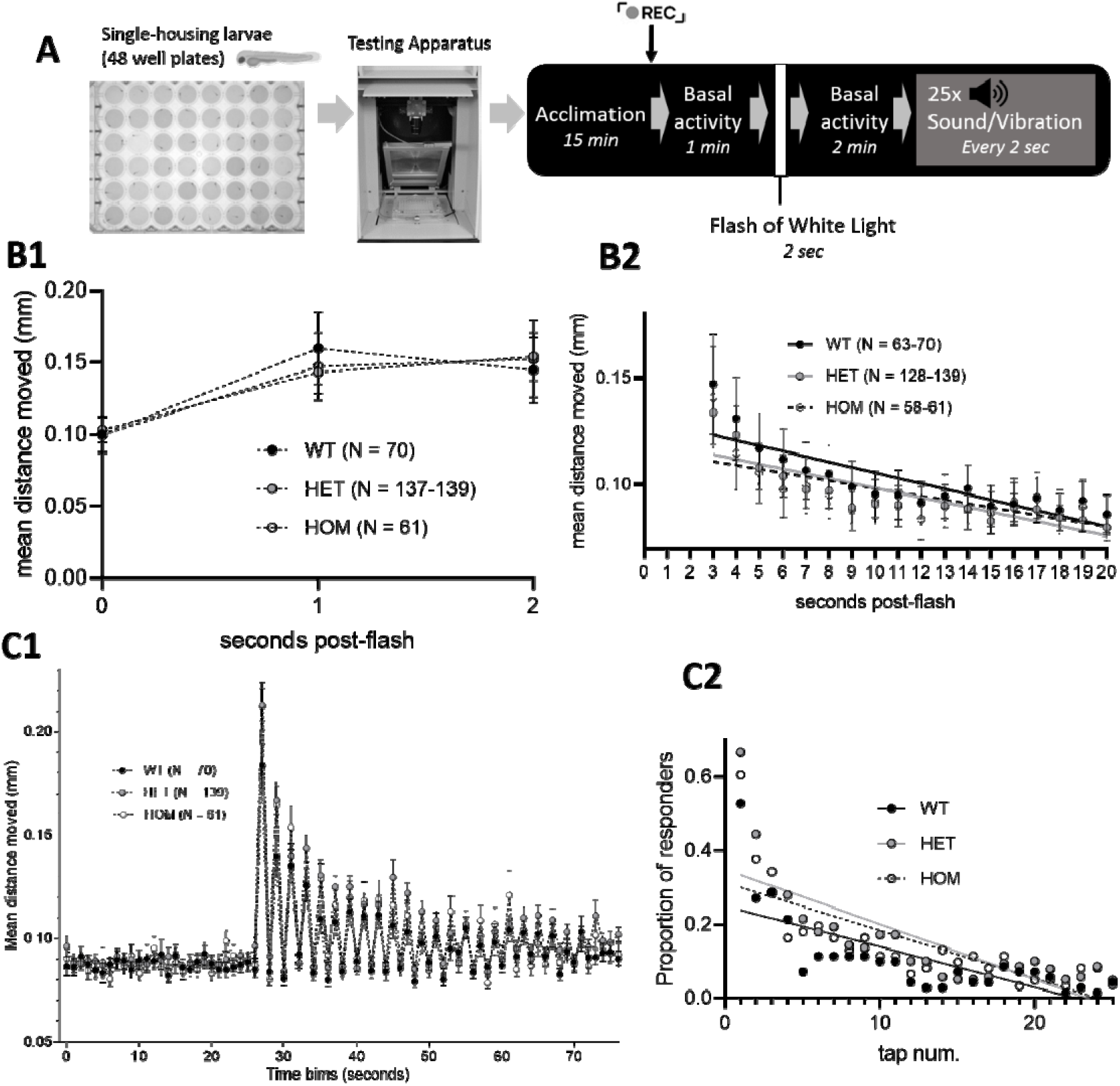
Loss of *baz1b* diminishes stress response and habituation in the flash of light and acoustic startle assay. **A)** Diagram of the assay. **B)** Flash of light: total distance moved during the flash of light (time 0) and two seconds after. Dots represent distance travelled in 1 sec time bins (B1), and rate of recovery for the 18 seconds following this period (B2). Dots represent 1 sec time bins. C) Acoustic startle assay: total distance moved 25 seconds before and during the acoustic cues (C1) and proportion of responders with calculated linear regression during the acoustic cues (C2). In all cases, data shows mean ± SEM. Intragroup variations in N number are due to statistical exclusion criteria applied at each time bin.

Next, we evaluated two parameters, distance travelled and proportion of responders, to assess the rate of habituation to a repeated acoustic startle. Defective startle habituation is associated with stress disorders (31). First, the total distance moved by fish during TAP events was used to define the slope of habituation and the significant effect of genotype-tap number interaction was found (F(2) = 8.42, *p* = 0.015) (Figure 4C1). Mutant fish habituated more quickly than WT, but the difference was significant only for HET fish and nearing significance for the HOM (HET: *p*= 0.011; HOM: *p*= 0.088). We also assessed the habituation response to repeated acoustic startle stimulation by looking at the decrease in the proportion of responders across stimuli (Figure 4C2). Statistical analysis showed a significant effect of genotype (F (2) = 29.283, *p*< 0.001). Post-hoc analysis revealed that in both HET and HOM the decrease in the proportion of responders was significantly steeper than WT siblings (Het: *p*< 0.001; HOM: *p*< 0.001).

The second line generated, *baz1b^ins35^*, shows similar tendency in the three behavioural assays (supplementary Figure 4).

### Reduced anxiety-like behaviour in *baz1b* LoF persists at later life stages

As both WS individuals and domesticated species show reduced stress-reactivity throughout life, we also investigated if the diminished stress-reactivity observed during early development in the *baz1b* LoF larvae was maintained at later stages. We used the novel tank diving assay, the zebrafish equivalent to the open field test in rodents (32), to assess anxiety-driven responses of the juvenile fish (10-12 weeks old, figure 5).

**Figure 5.**
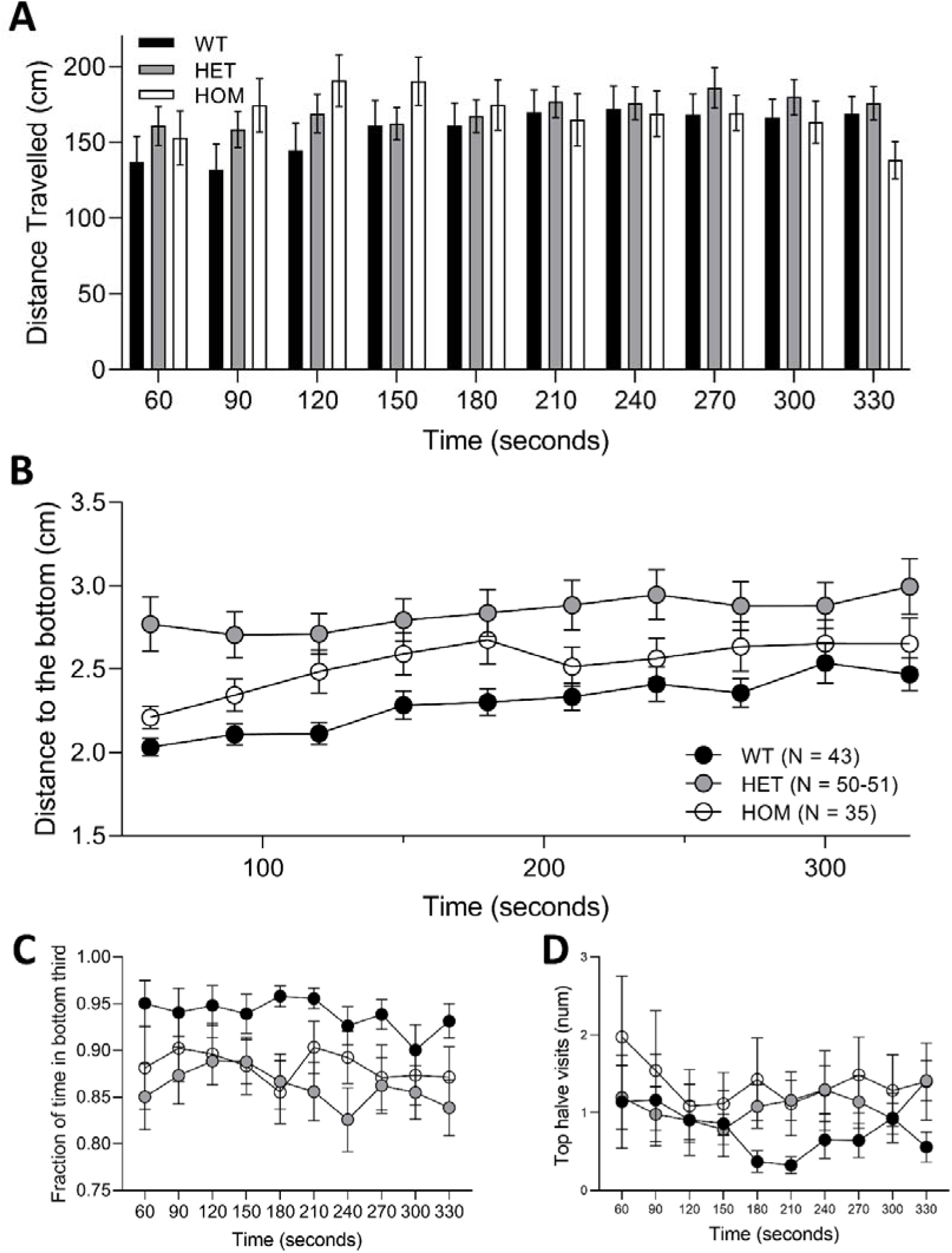
Novel tank diving assay shows anxiolytic phenotype in *baz1b* LoF zebrafish. **A)** Changes in distance travelled for the three genotypes in the novel tank diving assay divided in 30 seconds time beams. **B)** Distance to the bottom of the assay in each 30 seconds for the three genotypes. **C)** Fraction of time in the bottom third and **D)** top halve visits for the three genotypes during the novel tank diving assay. Graphs show mean ± SEM. Intragroup variations in N number are due to statistical exclusion criteria applied at each time bin.

There was a significant main effect of time (F (9) = 9.085, *p*< 0.0001) and genotype (F (2) = 7.336, *p* < 0.0001) on the distance to the bottom, but the interaction time-genotype failed to reach significance (F (18) = 1.029, *p* = 0.421). Distance from the bottom increased with time suggesting an anxiolytic tank diving response, which agrees with the normal response of fish to novelty. Genotype differences on the distance from the bottom were driven by HET fish spending significantly less time closer to the bottom than the other two genotypes (HET vs HOM: *p* = 0.0126; HET vs WT:*p* < 0.0001; HOM vs WT: *p* = 0.5475). Regarding total distance travelled, there were no significant effects of either genotype (F (2) = 1.579, *p* = 0.2116) or time (F (9) = 2.205, *p* = 0.1898).

When assessing the proportion of time spent on the bottom third of the tank (figure 5C), we found no effect of time (F (9) = 1.133, *p* = 0.3609), and no significant interaction between genotype and time (F (18) = 0.246, *p* = 0.9994). However, there was a significant effect of genotype (F (2) = 11.043, *p* < 0.0001). Post-hoc analysis revealed this to be driven by HET spending less time in the bottom third of the tank than WT (HET vs HOM: *p* = 0.0704; HET vs WT: *p* < 0.0001; HOM vs WT: *p* = 0.0793). In contrast, we did not find a significant effect of genotype on the visits to the top half of the tank (figure 5D; F (2) = 2.415, *p* = 0.12), while there was a significant effect of time (F (9) = 3.118, *p* = 0.0028) and the interaction between genotype and time (F (18) =2.7, *p* = 0.0001) whereby mutant fish are quicker to make more top half visits than wildtypes after an initial reduction observed in all three genotypes.

Again, both HET and HOM *baz1b^ιns35^* mutants showed similarly reduced anxiety-like behaviour although this did not reach significance (supplementary figure 5).

### Loss of *baz1b* disturbs the ontogeny of social behaviour in zebrafish

As the NCDS hypothesis predicts an extension of the socialization window during development (3, 4, 10, 33, 34), we assessed the ontogeny of social behaviour (35) in *baz1b* LoF mutants (figure 6 and supplementary figure 6).

**Figure 6.**
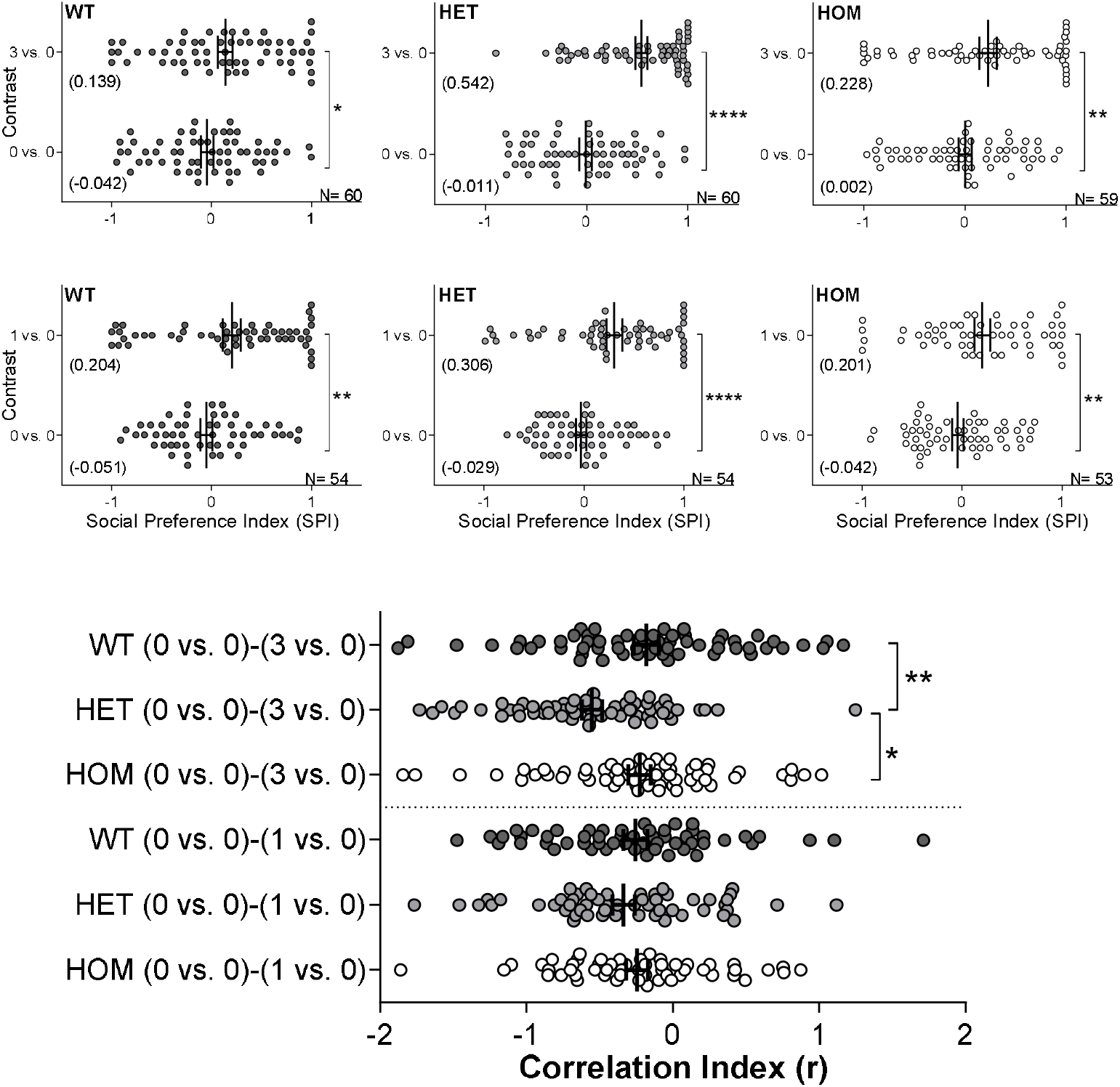
Loss of baz1b disturbs the ontogeny of sociability in zebrafish. Graphs show change in social preference index (SPI) between basal conditions (0 vs. 0) and contrast (0 vs. 3 or 0 vs. 1) for either WT, HET or HOM at week 2 of age (for week 1 and 3 see supplementary figure 6). N is included in each graph. Below panel shows the comparison of the correlation indexes for all the contrast at the corresponding week. A) week 1 (from 6 to 9 dpf), B) week 2 (from 13 to 15 dpf), C) week 3 (from 13 to 15 dpf). Graphs show individual values and mean (also between brackets for SPI) ± SEM. In all cases: * p < 0.05; ** p < 0.01; **** p < 0.001

Statistical analysis of the correlation indexes showed that there was a significant genotype-age of testing interaction on the social preference in the 0 vs. 3 contrast (F (4, 554) = 4.130, *p* = 0.0026). In contrast, the interaction did not reach significance in the 0 vs. 1 contrast (F (4, 556) = 0.06217, *p* = 0.9929).

None of the genotypes showed any significant preference for the social cues when presented with either contrast (0 vs. 3, or 0 vs. 1) during the first week of age and, consequently, no differences in the correlation indexes were detected (supplementary figure 6A).

At 2 weeks of age (figure 6), fish from all genotypes showed a significant tendency to remain in proximity to multiple conspecifics (0 vs. 3; WT: W = −544.0, *p* = 0.0451; HET: W = −1618, *p* < 0.0001; HOM: W = −770.0, *p* = 0.0032). Similarly, all genotypes showed this social preference when confronted with one conspecific (0 vs. 1; WT: W = −733.0, *p* = 0.0013; HET: −911.0, *p* < 0.0001; HOM: −699.0, *p* = 0.0016). Statistical analysis of the correlation index revealed that there was a significant difference between genotypes (H (6) = 17.22, p = 0.0041) and multiple comparisons revealed it to be driven by HET showing a greater motivation to interact with conspecifics than WT (*p* = 0.0048) and HOM (*p* = 0.0111) in the 0 vs. 3 contrast, while no differences were detected in the 0 vs. 1 contrast. There were no significant differences in the number of fish showing antisocial behaviour (avoiding arm with conspecific/s).

At 3 weeks of age (supplementary figure 6B), all genotypes showed a strong preference to interact with either multiple (0 vs. 3; WT: W = −1966, *p* < 0.0001; HET: W = −1596, *p* < 0.0001; HOM: W = −1924, *p* < 0.0001) or single (0 vs. 1; WT: W = −2286, *p* < 0.0001; HET: W = −2563, *p* < 0.0001; HOM: W = −2245, *p* < 0.0001) conspecifics and the statistical analysis of the correlation indexes did not describe any significant differences between genotypes (H (6)= 452, *p* = 0.1081).

Thus, results of the sociability assay suggest that *baz1b*-dosage influences the ontogeny of prosocial behaviours. Mutants of the *baz1b^ins35^* line also showed altered social ontogeny, although with variations: HOM showed increased motivation to shoal at week 1 but decrease at 3 weeks of age (supplementary figure 7).

## Discussion

We have shown that *baz1b*-deficient zebrafish recapitulate both morphological and behavioural phenotypes associated with the DS. In agreement with previous data, Baz1b seems to regulate the developing neural crest and its inputs to target tissues, thus resulting in both mild developmental delay at earlier stages and sustained morphological differences later in life. Lack of *baz1b* also resulted in fish showing a diminished stress response and increased willingness to socialise with conspecifics during the ontogeny of this behaviour.

Lack of functional *baz1b* resulted in minor morphological differences associated with mild developmental delay during early embryogenesis that resolved at later stages. The morphological delay observed at 12-15 hpf suggests that *baz1b* LoF might be influencing the beginning of neurulation (the transition from neural plate to neural tube) which in zebrafish embryos starts at 10 hpf, when the gastrulation is finishing. Similarly, morpholino-induced knockdown of *sox2*, a gene directly regulated by *baz1b* (12), in zebrafish show an aberrant pattern of neurulation at similar developmental stages which results in shorter body length as measured at 52 hpf (29). Although this mild developmental delay resolved at around 24 hpf, both changes in pigmentation and reduced rate of hatching were observed at later stages, which can be attributed to altered input of the neural crest to destination tissues: differential distribution of pigment cell within dermis and epidermis (36) and immature neuromuscular junction (37) respectively.

Given the observed morpho-developmental differences and the postulated role *baz1b* plays in neural crest (11, 12), we also assessed the expression pattern of different neural crest markers by *in situ* hybridization. We observed an increased pattern of expression for *crestin* and a reduction in *pax2a/b*, *sox2* and *epha4*, which suggest that lack of *baz1b* shifted the balance between pre-migratory and migratory neural crest cells consistent with an extended immaturity period of the neural crest. In contrast, no changes in *foxd3* or *sox10* were observed indicating that, although *baz1b* LoF alters the maturation of the neural crest, it does not seem to affect neural crest formation and maintenance. Thus, lack of *baz1b* leads to mild neurocristopathy, which is in line with previous findings (11, 12) and points at *baz1b* as a major regulator of neural crest development.

We also assessed the expression pattern of *robo1/2* and *foxp2*, genes the expression of which is regulated by Baz1b and are associated with neurological features relevant to DS such as vocalization (11). Interestingly, there was no detectable difference in the expression of these three genes, which could be explained by the lack of such vocal communication in zebrafish. Nonetheless, the expression pattern of both *robo1* and *foxp2* becomes neuroanatomically and functionally more complex during zebrafish adulthood (38, 39). Thus, Baz1b could still be influencing their expression in zebrafish but at later stages to the ones studied here.

Deficiencies in *baz1b* also induced mild differences in cranial bones and facial features at both 5 dpf and 30 dpf but did not affect the external maturity of the fish. In the main line of this study, *baz1b^del44^*, the LoF led to a significant elongation between Mechkel’s cartilage and the ceratohyal bone and, consequently, an increase in the eye-snout distance in the heterozygous larvae. On the contrary, the differences in craniofacial features reported in the second line, *baz1b^ins35^*, were associated with shortened snouts, which is more consistent with the reduction in craniofacial features associated with the DS. Nonetheless, the difference in phenotypes seen in the two *baz1b* LoF lines suggests that, although *baz1b*-dosage plays a role in the determination of facial features, other genes might be influencing the overall phenotype. This might explain why the reduction in facial features is sporadically reported in different species of domesticated animals (6–8). Recent findings comparing facial patterns in rats based on their levels of tameness and aggression have reached a similar conclusion (40).

Behaviourally, deficiencies in the levels of functional Baz1b led to a reduction of stress reactivity at 5 dpf. The results from the three behavioural assays at which fish were exposed (FLDT, quick flash of light and repeated acoustic startle) suggest that *baz1b* LoF zebrafish habituate more quickly to both light and acoustic perturbations. Although differences in basal activity were reported in the FLDT assay, those were not observed in the other assays. The quick flash of light assay is an adaptation of that recently developed by Lee and colleagues (41) which is shown to rely on HPI reactivity, the equivalent to the mammalian HPA axis. Therefore, our data are consistent with reduced HPI response and suggest that maturity of the HPI axis is affected by the deficits of Baz1b during development in agreement with the NCDS hypothesis.

The reduced stress reactivity observed at 5 dpf is maintained later in life in the *baz1b* LoF zebrafish. The tank diving test showed that, although all fish developed an anxiety-like behaviour that gradually diminished throughout the time spent in the assay, heterozygous fish showed significantly reduced anxiety-like behaviour when compared to WT. Therefore, our data suggest that deficits in *baz1b* diminished anxiety-related responses in juvenile zebrafish in agreement with the NCDS hypothesis.

The sociability assay suggest that *baz1b*-dosage influences the ontogeny of pro-social behaviours, as observed in domesticated species. Our data showed that *baz1b* LoF resulted in an increased inclination to interact with conspecifics at 2 weeks of age. Furthermore, the sociability assay also shows that multiple, instead of single, conspecifics are stronger drivers of social behaviours in *baz1b* LoF zebrafish, which agrees with the NCDS hypothesis as this alteration will be favouring larger social groups and intraspecific interactions (5, 9). Although *baz1b* LoF might not be influencing the overall time it takes to develop zebrafish’ social skills, it seems to affect the eagerness to establish social contact with conspecifics during its ontogeny.

Differences in sociability are no longer observed by 3 weeks of age. However, the lack of differences could be a consequence of either: a) all fish, regardless of genotype, have developed a similar level of sociability at this age thus suggesting differences observed at early stages have resolved; or b) if differences in sociability remain, they cannot be detected in this assay as the salience to socialise is too strong by 3 weeks of age in all genotypes. Nonetheless, Baz1b might still be playing a role in social responses later in life as it has been implicated in stimulus-specific response to different emotional paradigms in rodents (42).

Taken all together, our research supports the NCDS hypothesis and points at *baz1b* as one of the main genes regulating both morphological and behavioural DS-associated changes in zebrafish. This investigation also supports zebrafish as a feasible vertebrate model to assess the effects of gene dosage-variation in both morphological and behavioural endophenotypes of domestication and human self-domestication-associated disorders (5, 13).

Although both lines used in this study revealed similar findings, several differences arose. Within the *baz1b^del44^* line, only heterozygous zebrafish showed both morphological and behavioural changes when compared to WT while homozygous remained mostly unchanged. On the other hand, the *baz1b^ins35^* line showed similar changes in both heterozygous and homozygous but less marked. This incongruency could be manifesting different penetrance of the mutation since the fold change of *baz1b* expression is minor in the second line. Nevertheless, Baz1b is suggested to influence these characteristics in a dosage manner while its total absence leads to mortality (12, 43). Its haploinsufficiency, as seen in WS, is associated with hyper-sociability (13, 14) while its duplication is linked to opposing behaviours such as autism-like features (15, 16). Therefore, heterozygous individuals might reflect better both the human pathology and the gene copy-number variations observed in domesticated species.

Finally, Baz1b, which acts as both a transcription factor and chromatin remodeller in the neural crest, could be in a prominent position to extend the comparative studies to the epigenome. Targeting Baz1b-associated chromatin interactions by novel multi-omics approaches might help to elucidate the associated changes in gene expression. Some species of recently domesticated vertebrates show epimutations rather than gene mutations (44). Interestingly, most of those epimutations are directed to the same set of genes, including most of those expressed in the neural crest, that are observed in established domesticates. Thus, future studies that investigate how Baz1b regulates the epigenomic landscape can help to elucidate the set of regulatory changes leading towards the initial domestication-associated phenotypes.

## Materials and methods

### Zebrafish husbandry

All *in vivo* experimental procedures described were reviewed and approved by the QMUL ethics committee (AWERB) following consultation of the ARRIVE guidelines (NC3Rs, UK) and conducted in accordance with the Animals (Scientific Procedures) Act, 1986 and Home Office Licenses.

Zebrafish were housed in a recirculating system (Techniplast, UK) with a light:dark cycle of 14:10. Housing and testing rooms were maintained at ~25–28 °C. Subjects were maintained in aquarium-treated water and fed twice daily with dry food (ZM-400,Zebrafish Management Ltd, Winchester, United Kingdom) in the morning and live brine shrimp (*Artemia salina*) in the afternoon. All zebrafish used from this study originated from a Tubingen wild type (WT) background line.

To breed them, zebrafish were moved to breeding tanks with perforated floors in the evening and eggs collected the following morning. Eggs were incubated in groups of no more than 50 per Petri dish at 28 °C until 5 dpf. At 6 dpf, larvae were transferred to the recirculating system and fed twice daily with commercial fry food (ZM-75, ZM-100, Zebrafish Management Ltd, Winchester, United Kingdom) and life paramecium/brine shrimp, depending on their age.

### Generation of *baz1b* loss-of-function (LoF) zebrafish stable lines using CRISPR/cas9

The procedure used is similar to that described by Keatinge and colleagues (45) but with small variations. One-cell stage zebrafish embryos were injected into the yolk with 1 nL injection solution containing 62.5 ng/μL crispr RNA (crRNA, Sigma), 62.5 ng/μL tracrRNA (Sigma, cat.TRACRRNA05N), 1:8 dilution Cas9 protein (New England Biolabs, cat. M0386M; diluted in buffer B, New England Biolabs, cat. B802S) and 1:40 dilution of phenol red (Sigma Aldrich). The crRNA (5’CUCAUCCUCCACCACCCAGG) was designated to target a section of exon 5 of the zebrafish gene *baz1b* (ensembl gene ID: ENSDART00000158503.2) overlapping a restriction site for the Bsl1 enzyme (New England Bioscience) which includes a PAM site. Once a pair of founders was identified (outcrossing them to WT and genotyping the offspring by PCR) they were in-crossed and resulting F1 genotyped. All identified homozygous individuals (*baz1b^-/-^*) from the F1 were outcrossed with multiple WT to obtain a F2 of heterozygous (*baz1b*^+/−^). Finally, F2s were in-crossed and F3 genotyped to establish a breeding stock of fish from each genotype. Primers used for genotyping: forward 5’ AGAAGAAGAAATGGGTCATGCC and reverse 5’ CCTCTTAAACCATCTCACCTTGT.

Zebrafish lines can be provided upon email request to the corresponding authors.

### Quantitative PCR (qPCR) to validate *baz1b* LoF lines

Groups of 16 larvae at 5 dpf from all corresponding genotypes (N = 3 per genotype) were collected and stored in RNAlater (Thermo Fisher) until use. Collected larvae were previously scrutinized to ensure consistent developmental stages across groups. RNA extraction and quantitative PCR (qPCR) were carried similarly as previously described (32). The relative expression of *baz1b* (Forward: 5’ CAGCAGTCAGCTCACGCATAAG; Reverse: 5’ TGCACAATCTCCAGAACAGGC; Primer efficiency: 96%; Product length: 111 bp of the mRNA spanning exon 1 and 2, upstream of the mutation site) was calculated with two housekeeping genes (*β-actin* and *rpll3α*) as references, adjusted by their corresponding efficiencies and normalised similarly as done before (32).

### Whole body *in situ* hybridization (WISH)

We used a standard method for *in situ* hybridizations (46). All embryos of 26 hpf or older were raised in fish water containing 200 μM 1-Phenyl-2-thiourea (PTU; Sigma Aldrich) to prevent melanin synthesis. A minimum of 15 individuals per genotype and RNA probe were carefully staged before WISH using Kimmel’s criteria (47) to ensure potential differences were due to genotype and not developmental delay. RNA probes were generated and amplified by PCR from cDNA produced similarly as above from WT larval tissue. PCR products were isolated with a QIAquick Gel Extraction kit (Quiagen) and purified fragments cloned into pGEM-T Easy vector (Promega). Specific details for the generated probes are as follow (gene name (accession number), forward primer (5’-3’), reverse primer (5’-3’), size in bp): *baz1b* (NM_001347672.1), GAAGGAGCCACAGTCCAAAGAG, TGACACAGAGCCAACAGCAGAC, 1728; *crestin* (NM_130993.2), AAGCGGATCGGAAGTGGGAAAG, CAGGCAGAAGCAGAGAGAACAG, 1235; *foxd3* (NM_131290.2), GGAAAAGAGACCCAAGCAAGCC, ATATTTGACGGGACGCTGAGG, 1200; *foxp2* (NM_001030082.2), ACAGCACCCAGGCAAACAAG, GGATTACCCAGAAGCGGCAAAG, 1320; *sox2*(NM_213118.1); CATCAAGAGACCCATGAACGCC, TCGTGCCGTTAATCGTCGTACC, 818; *sox10*(NM_131875.1), TGGCGGGAAAACGCAGATAAAG, ACCACAACAGCACACGACAG, 1618; *pax2a* (XM_017358685.2), AAACAATCCATCTCCCCCTCCC, GCTGAGGCCATGTTGCTTTTTG, 1046; *robo1* (NM_001365166.1), GACGACAGTTACGACACAGACC, ATTCTTGCCCGTTCTCTGACCC, 1131; *robo2* (XM_021481460.1), TCTCCGCCCTAACACCATCTAC, CCACCTTTTCCACCTGACAGAC, 1539. cDNA clones for the following probes were obtained from others: *epha4* (48) and *pax2b* (49). Pictures were taken using a Leica S9i stereo microscope with an integrated CMOS camera (10Mpixels, pixel size 1.67 μm x 1.67 μm), in a laptop Dell Latitude E5440 with the software Leica Application Suite LAZ EZ version 3.4.0 and stored as tagged image file format (tiff).

### Alcian blue staining and morphological measurements

5 dpf larvae were fixed overnight in 4% paraformaldehyde (PFA) in a phosphate-buffered solution with 0.1% Tween-20 (PBT; Sigma-Aldrich). Following day, larvae were washed with PBT, depigmented with a PBT solution containing 30% H2O2/0.5% KOH for approximately 1 hour, washed further with PBT, stained in 0.37% HCl/70% ethanol /0.1% Alcian Blue in PBT for 90 minutes and washed again with 1% HCl/70% ethanol in PBT. Larvae were then cleared with 50% glycerol/0.5% KOH and pictures taken (same setup as above) from the ventral and lateral planes and analysed using software ImageJ Java 1.8. 0_45 [64-bit] (Rasband, W.S., ImageJ, U. S. National Institutes of Health, Bethesda, Maryland, USA, https://imagej.nih.gov/ij/, 1997-2018). Bone measurements taken: 1) width of the ceratohyal (CH) at palatooquadrate (PQ) joint; 2) length of (1) to Mechkel’s cartilage (M); 3) length M-CH; 4) width M at PQ joint; 5) M-PQ angle; and 6) CH angle.

### Additional morpho-physiological measurements

Multiple individual breeding pairs were set to breed simultaneously to obtain fish from the three possible genotypes (WT x WT, HOM x HOM and WT x HOM). 30 minutes after breeding, eggs were collected, pooled together according to genotype and 150 eggs each selected for morphophysiological analysis (3 plates each). Pictures were taken at different stages using a Leica TL3000 Ergo microscope with a Leica DFC3000 G camera, the software Leica Application Suite X version 3.4.2.18368 and collected as tiff. Following morphological analysis were performed using ImageJ: head-to-tail distance at 12 and 15 hpf (50), whole body length (BL) and portion of tail protruding from yolk sack at 24 hpf, relative pigmentation intensity at 30 hpf (50), and BL at 60 and 72 hpf. Additionally, the percentage of hatched larvae was monitored between 49 to 73 hpf (50).

At 5 dpf, some larvae (N = 30 per genotype) were fixed overnight with PFA. Following morning, larvae were briefly washed with PBST and cleared with increasing concentrations of glycerol up to 100%. Whole body dorsal and lateral plane pictures were taken using the same setup as in the Alcian blue section to measure similar parameters as assessed in Martinez et al., 2019 (51): BL, eye length (EL), eye-snout distance (ESD), eye width (EW), head-trunk angle (HTA), head width (HW) and inter-ocular distance (IOD). 30 dpf fish were processed likewise, but without clearing with glycerol, to measure BL, EL, ESD, EW, HTA, HW and IOD, and to assess their maturity state based on pigmentation/maturation patterns similarly as Singleman & Holtzman 2014 (30).

To ensure observed differences were not due to angle variations of the samples when taking pictures, fixed fish were positioned on a grooved area carved out of a 1% agarose base at the bottom of the Petri dishes.

### Behavioural evaluation

Larvae used for behavioural assays were obtained from in-crosses of *baz1b* heterozygous (HET x HET) zebrafish and thus examiner was blind to genotype until larvae were culled and genotyped by PCR (similar to above) after behavioural experimentation.

#### Forced light-dark transition (FLDT) at 5 dpf

This test was performed between 9 am and 4 pm. 5 dpf larvae were individually placed in 48-well plates and set to acclimate for 15 minutes inside the dark DanioVision Observation Chamber (Noldus Information Technology, Wageningen, The Netherlands). After this period, an initial 10-minute recording in the dark (infrared conditions) was used as baseline and zebrafish larvae were subjected to 3 consecutive forced light/dark transition cycles, each consisting of 10 minutes of light followed by 10 minutes of dark. Distance travelled was recorded using Ethovision XT software (Noldus Information Technology, Wageningen, NL) and data were outputted in 10-second time bins. To account for experimental variation, two independent experiments per day were conducted on three consecutive days.

#### Flash of light and acoustic startle habituation at 5 dpf

Sensorimotor startle response and habituation were tested in a protocol combining a variation of the light-locomotor behavioural assay developed by Lee and colleagues (41) and a startle tap test. This assay was performed between 9 am and 4 pm. At the beginning of each trial, individual 5 dpf larvae resulting from *baz1b* heterozygous in-crosses (unknown genotypes until the end of the experiment) were randomly placed in a 48-well plate and set to acclimate inside a dark DanioVision Chamber. After 10 minutes of initial habituation, recording started. Following a further minute of acclimation in the dark, larval zebrafish were exposed to a brief illumination in white light (2 seconds), and then back to dark conditions for 2 minutes. Right after, zebrafish larvae were subjected to 25 sound/vibration consecutive stimuli with an inter-stimulus interval of 2 seconds. Distance travelled was recorded using Ethovision XT software and data were outputted in one-second time bins. Two independent experiments per day were conducted in three consecutive days.

#### Novel tank diving assay of juvenile zebrafish

This assay was performed as described previously (32) with small adaptation for juvenile fish. Zebrafish were housed in tanks according to genotype. At least two batches of fish were used per genotype (to avoid tank effect) at 10-12 weeks old. This assay was conducted between 9 am and 2 pm. Trapezoid tank dimensions were 22 (bottom part) x 27 (top part) x 9 × 14.5 cm. It contained 2 litres of fresh fish system water (10 cm height water column) that was changed within trials. Fish were individually placed in the test tank and recorded from the side for 6 minutes. Swimming activity, including distance to the bottom and total motility, were tracked and analysed using EthoVision software and data outputted in 30-seconds time bins.

#### Ontogeny of sociability

This assay was performed as Dreosti et al. 2015 (35) with small modifications. Briefly, 4 tanks of 50 fish were kept for each genotype and re-analysed at weeks one (6 to 8 dpf), two (13 to 15 dpf) and three (20 to 22 dpf). Tests were performed between 10 am and 7 pm. Each fish was only tested once within each week, with a delay between consecutive weeks of testing of ≥ 7 days. Fish used as test subjects from each genotype were age/size matched to stimuli fish from the same genotype (same housing tank). Experiments were performed simultaneously in two DanioVision Observation Chambers, each containing 6 arena setups with fish from a certain genotype. At least two independent experiments were conducted each day per genotype and DanioVision Chamber with side of social cue presentation balanced in each trial. Swimming activity, including total motility and position within the arena, were tracked and analysed using EthoVision XT software and data outputted in single 15 minutes bins. Social Preference Index (SPI) was calculated similarly to Dreosti et al. 2015 but, additionally, we calculated the Correlation Index (r) to compare predisposition of fish to socialize: [r = SPI_ExperimentalPhase_ – SPI_AcclimationPeriod_]

### Statistics

Power calculations (beta = 0.8, alpha = 0.05), with effect sizes determined from pilot studies, were used to estimate sample size for each experiment. Whenever possible, animals/contrast/testing order were selected at random. Unless stated otherwise, all effects are reported as significant at *p* < 0.05.

#### Molecular and morphological comparisons

The qPCR data on relative change in gene expression and all morphological comparisons were analysed using GraphPad Prism 9.0.2 for Windows (GraphPad Software, San Diego, California USA, www.graphpad.com). Dataset was assessed for normal distribution by Shapiro-Wilk’s test (*p* > 0.05) and visually assessing the normal QQ plot. If normality assumptions were not violated, datasets were analysed with an ordinary one-way ANOVA with Tukey correction to account for multiple comparisons. If normality was violated, data was either transformed to log 10 or using other normalising factors (e.g. BL), or analysed using the nonparametric Kruskal-Wallis test for unpaired samples and using Dunn’s test to correct for multiple comparisons. Rate of hatching was analysed with a two-way repeated measures ANOVA with “genotype” and “time” as factors and “subject” as matched set and using Tukey correction to account for multiple comparisons.

#### Larval behavioural assays and novel tank diving

Corresponding data was analysed using R version 4.0.0 and Rstudio version 1.2.5042 and results of all statistical analyses were reported with respect to a type-1 error rate of α = 0.05. Data analysis was performed similarly as previously described (52) but with small changes.

For FLDT, we created three subsets of data to be analysed separately: baseline, light, and dark periods. Baseline period was fitted to the linear mixed model with the total distance travelled as a response variable, genotype as fixed effects, and fish ID as random effects. Light and dark periods constituted the three similar events combined. To explore the change in larvae movement over time in the light (increase) and in the dark (decrease) over time, linear models at light and dark periods were fitted using distance travelled as response variable, interaction between genotype and time as independent variable and fish ID as random effects. Linear mixed models were calculated using R package lme4 and significant fixed effects identified using chi-squared test. To further characterise the effects, where significant differences were established, and post-hoc Tukey test using R package ‘multcomp’ were carried out. The β coefficient in light and dark period models represents the increase or decrease in distance travelled over time and can be interpreted as the larval ‘recovery rate’.

For the flash of light and acoustic startle habituation test, data was divided into 4 parts: baseline period 1, flash of light+recovery, baseline period 2 and response to startle stimuli. Baseline period 1 and 2 were analysed as described above. Response to the flash of light was analysed by looking at total distance moved in 2 seconds following the flash and the rate of recovery. In the “jump” analysis, data was fitted to the linear mixed model with the total distance travelled as a response variable, genotype as fixed effects, and fish ID as random effects. The slope was analysed as previously described in FLD analysis, where the linear model was fitted with distance travelled as response variable, interaction between genotype and time as independent variable and fish ID as random effects.

Response to startle stimuli was analysed in two different ways. In both approaches, each TAP event was defined as a two second event, consisting of exact time of the startle stimuli and the following second. In the first approach, we calculated slope of habituation to startle stimuli by fitting a linear mixed model using distance travelled as response variable, interaction between genotype and tap event as independent variable and fish ID as random effects. Then, significant fixed effects were identified using chi-squared test and, when significant differences were established, post-hoc Tukey test was used to further characterise the effects. In the second approach, we defined a response/non-response status for each fish with threshold for responsive status defined as mean distance moved per second during the basal period plus two standard deviations (SD). The threshold was calculated for all three genotypes together, as genotype was not a significant predictor of basal distance travelled. Each fish was assigned as ‘responder’ if it moved more than the threshold during the TAP event or as ‘non-responder’ if it did not. Using the R package ‘betareg’, we modelled beta regression with percentage of fish responding to stimulus as a response variable and interaction between TAP event number and genotype as explanatory variables. Then, likelihood ratio tests for nested regression models were performed to assess if the interaction between TAP event number and genotype was a significant predictor of individual responsiveness.

In the novel tank diving assay, data on distance travelled in the tank and distance from the bottom of the tank was fitted to a linear mixed model with the total distance travelled and distance from bottom in 30 second bins as a response variable, time and genotype as fixed effects, and fish ID as random effects. To analyse differences in the time spent in the bottom third of the tank, we performed beta regressions using the R package ‘betareg’. To analyse genotype differences in the number of transitions between the top and the bottom of the tank, we fitted the data to a generalized linear mixed model with Poisson distribution using the number of transitions to the top-bottom of the tank as response variable, time and genotype as fixed effects, and fish ID as random effects. In all novel tank diving data analyses, the first 30 seconds were excluded as recording of the assay is started prior to the fish being added.

#### Ontogeny of sociability behaviour

Corresponding data was analysed using GraphPad Prism 9.0.2 for Windows. An ordinary 2-way ANOVA was used to assess genotype x age interaction of the correlation indexes for each contrast (0 vs. 3 or 0 vs. 1). The differences in SPI each week of testing within genotypes were analysed similarly as Dreosti et al., 2015 (35): the non-parametric two-tailed Wilcoxon signed-rank test of paired samples. Correlation indexes were analysed with the nonparametric test of Kruskal-Wallis of not paired samples and using Dunn’s test to correct for multiple comparisons.

## Data availability

All data and statistics used will be available at https://github.com/Jovitope/baz1b-zebrafish.git(Publicly available once accepted).

## Author Contributions

JVTP: conceived and planned experiments, lines generation, molecular and behavioural experiments, data analysis and writing. SA: morphological measurements and analysis. AMM: analysis of 5 dpf behavioural data. WH: analysis of the novel tank data and part of the 5 dpf behavioural data. JGG: assistance in line generation. SEF and GV: provided funding. CHB: provided funding, supervised the project and writing. All authors discussed the results and contributed to the final manuscript.

## Acknowledgments

We thank Dr Katharine E. Lewis for providing the *pax2b* probe.

Funders: Human Frontiers Scientific Program (RGP 0008/2017, JVTP, CHB); National Institute of Health (NIH U01 DA044400-03, AMM, WH, CHB); Leverhulme Trust (RPG-2016-143, SA, CHB); European Commission (SPANUMBRA – 833504, SA, CHB); Ministerio de Universidades (Next Generation EU), Universitat de València, Ayudas María Zambrano para atraer talentos del ámbito internacional (ZA21-026, JVTP).

