## Supplementary Figure for "Behavioural analysis of loss of function zebrafish supports *baz1b* as master regulator of domestication"

Supplementary figure 1. **Additional WISH showing *baz1b^del44^* LoF induces mild neurocristopathy in zebrafish**. **A)** *baz1b*, **B)** *sox10*, **C)** *foxd3*, **D)** *epha4*, **E)** *robo1*, **F)** *robo2*, **G)** *foxp2*. In all figures, WT, HET and HOM are represented and scale bar depicts 2 µm. Corresponding hpf are shown in each panel.

Supplementary figure 2. **Deficits in *baz1b^ins35^* lead to mild neurocristopathy in zebrafish. A)** Left panel shows a DNA blast for the portion of *baz1b*’s exon 5 amplified by PCR used for genotyping between WT (above) and mutants (below, 35 bp insertion). Primers used are highlighted in yellow, crRNA site in blue, PAM site in grey and restriction enzyme over-lined. Right panel shows the in-frame translation to amino acids (aa) sequences in WT (above) and mutants (below). Changed aa are in red and stop codon marked as a dash (-). **B)** Relative change in gene expression (log_10_) assessed by qPCR between 5 dpf larvae from each genotype. Figure shows individual values (N = 3 per genotype, each representing a group of 16 larvae combined) and mean ± standard error mean (SEM). **C)** Whole larvae *in situ* hybridization (WISH) against *crestin* for WT, HET and HOM at 18 and 24 hpf. Scale bar represents 2 µm. **D)** Similar WISH than C) but against *pax2a*, *pax2b* and *sox2*, genes regulated by *baz1b*. **E)** Developmental comparison between the three phenotypes for the head to tail distance at 12 hpf and at 15 hpf, ratio body/tail-out-of-yolk-ball at 24 hpf, relative percentage of pigmentation at 30 hpf, body length at 60 hpf and at 72 hpf, and percentage of hatched larvae between 49 to 73 hpf. Graphs show mean ± SEM. In all cases: * *p* < 0.05; ** *p* < 0.01; *** *p* < 0.001.

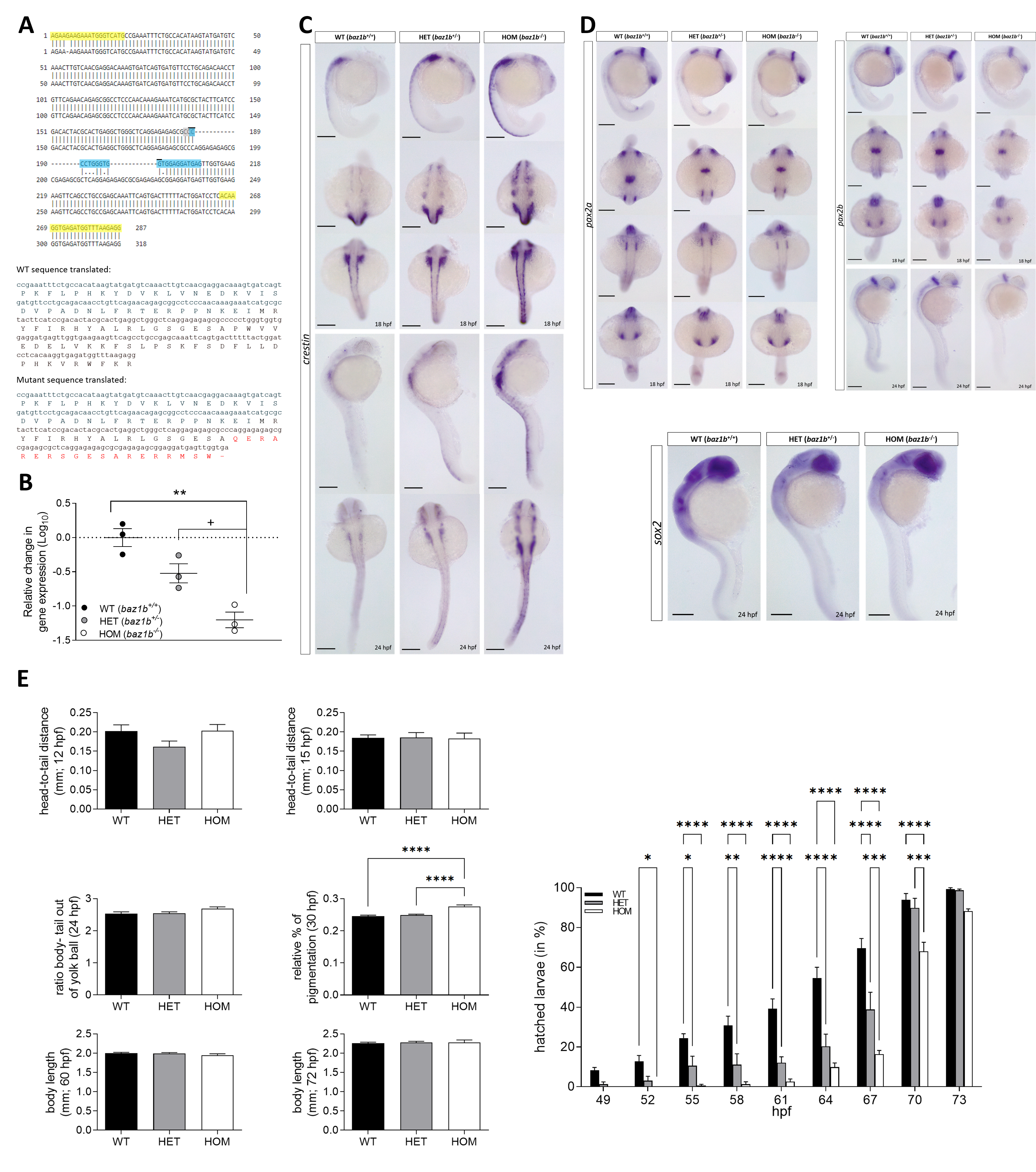

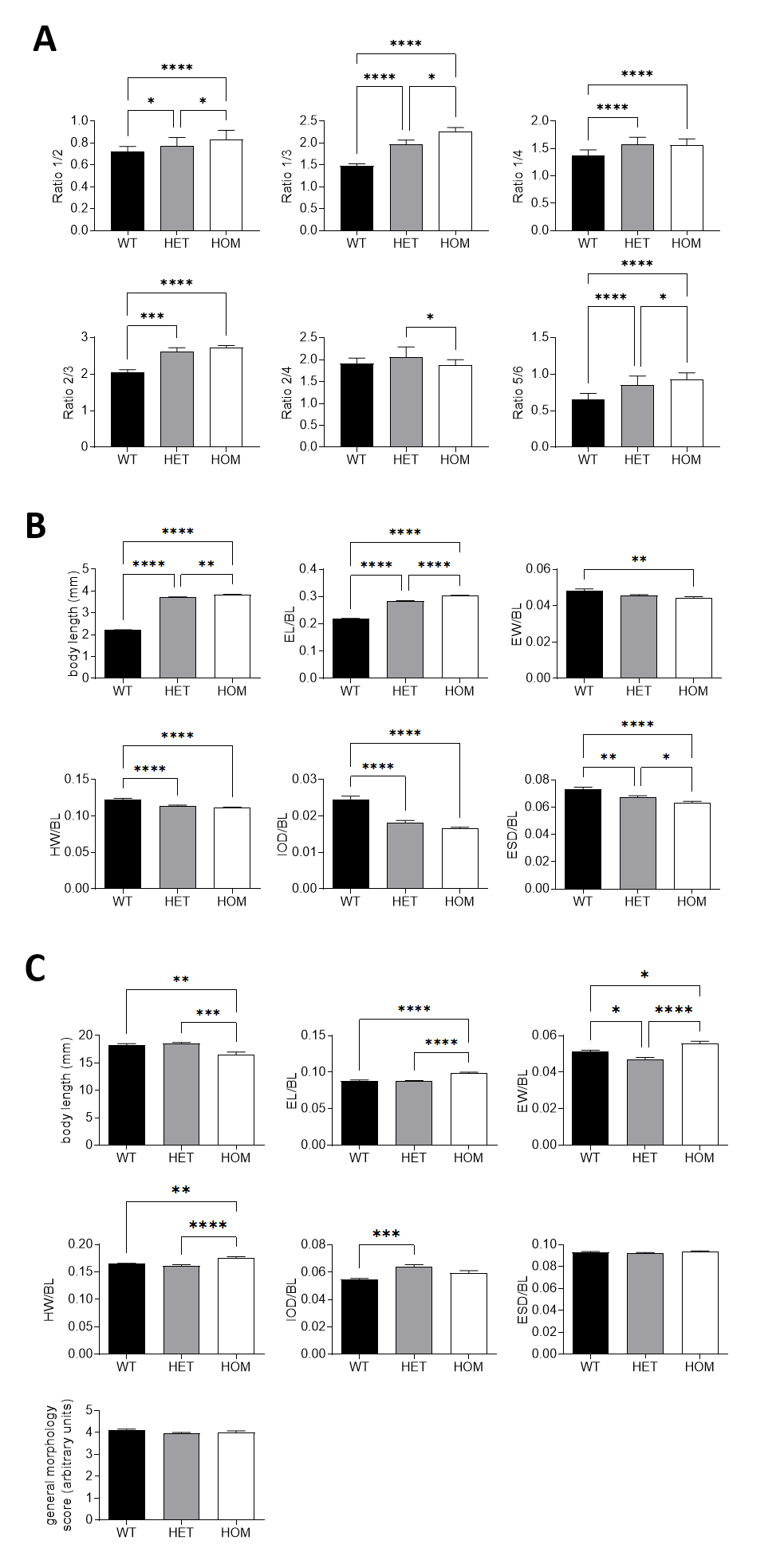

Supplementary figure 3. ***baz1b^ins35^* LoF zebrafish have altered craniofacial features**. **A)** Comparison between genotypes for the cranial features observed at 5 dpf by alcian blue staining: ratio 1/2 (A1), ratio 1/3 (A2), ratio 1/4 (A3), ratio 2/3 (A4), ratio 2/4 (A5) and ratio 5/6 (A6). **B)** Morphological comparisons of intact 5 dpf for the three genotypes: body length (BL, in mm; B1), eye length (EL) normalised to BL (B2), eye width (EW) normalised to BL (B3), head width (HW) normalised to BL (B4), inter-ocular distance (IOD) normalised to BL (B5) and eye-snout distance (ESD) normalised to BL (B6). **C)** Similar morphological comparison than B) of intact juvenile fish (30 dpf) and the general morphology score to assess maduration (C7). Graphs show mean ± SEM. In all cases: * *p* < 0.05; ** *p* < 0.01; *** *p* = 0.001; **** *p* < 0.001

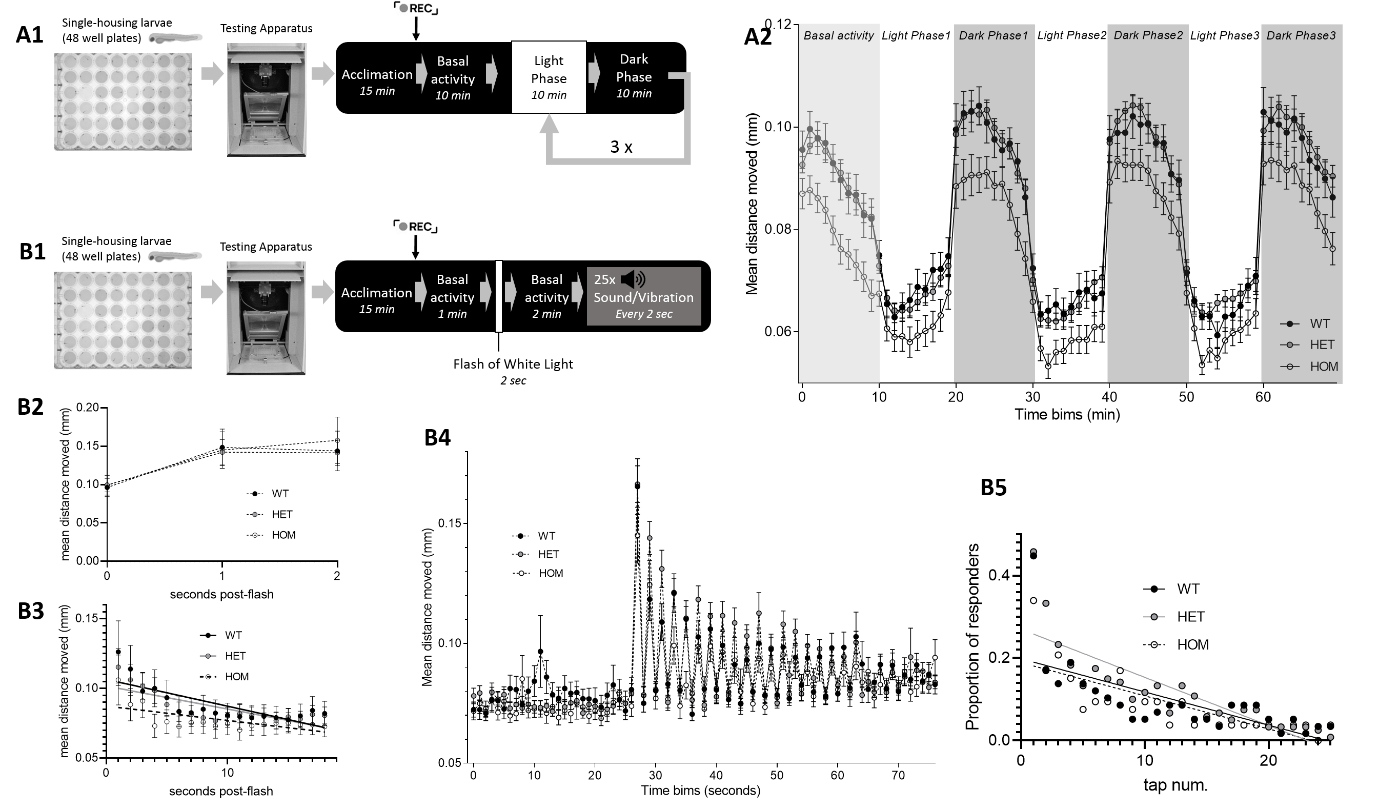

Supplementary figure 4. **Loss of *baz1b^ins35^* diminishes stress response and habituation in larval zebrafish**. **A)** Forced light dark transition (FLDT) assay: diagram of the assay (A1) and mean distance moved per genotype during the baseline and the three light-dark cycles (10 minutes each). **B)** Flash of light and acoustic startle habituation assay: diagram of the assay (B1); total distance moved during the flash of light and two seconds after (B2) and rate of recovery for the 18 seconds following this period (B3); total distance moved 25 seconds before and during the acoustic cues (B4) and proportion of responders with calculated linear regression during the acoustic cues (B5). In all cases, data shows mean ± SEM.

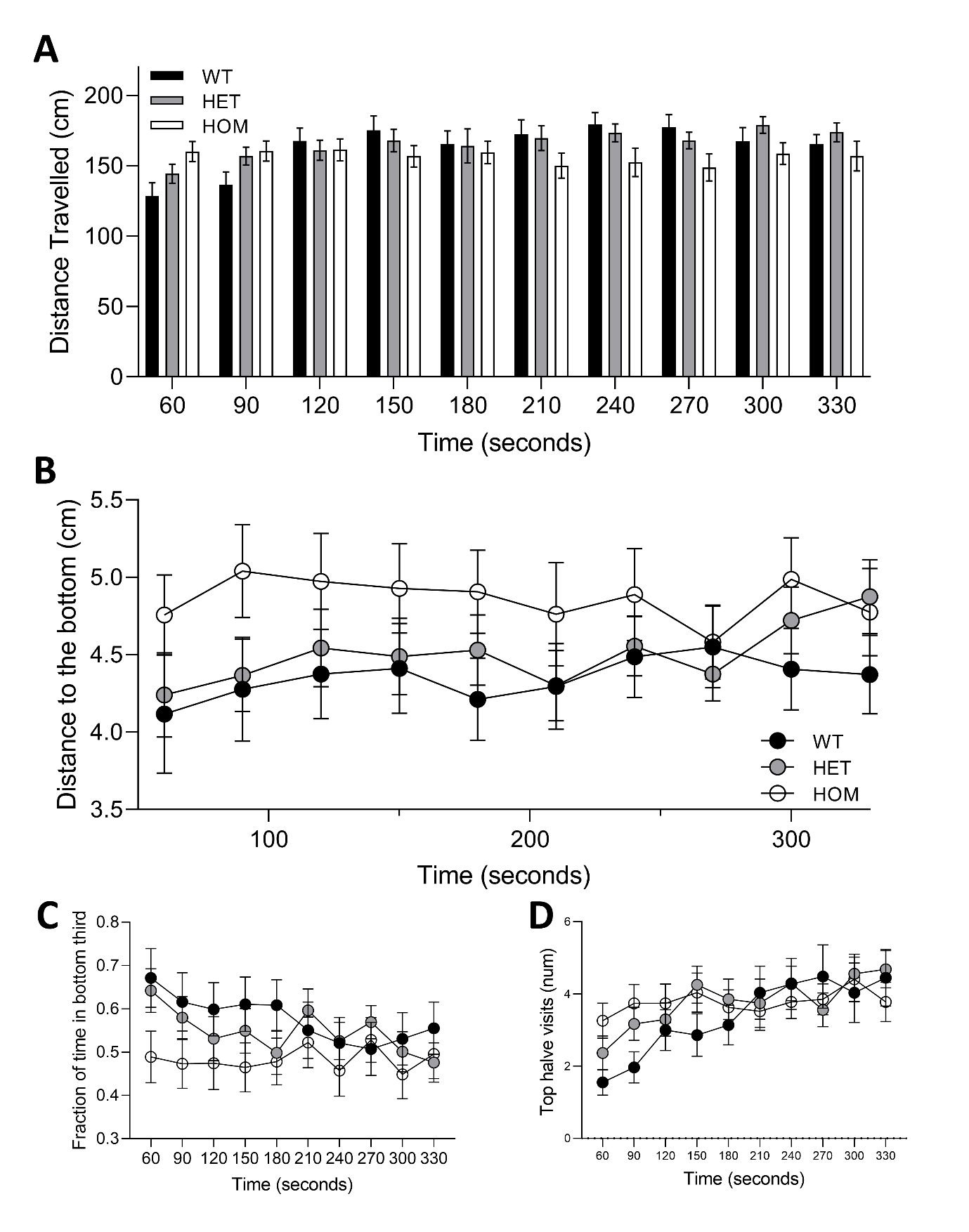

Supplementary figure 5. **Novel tank diving assay shows anxiolytic phenotype in *baz1b^ins35^* LoF zebrafish**. **A)** Changes in distance travelled for the three genotypes in the novel tank diving assay divided in 30 seconds time beams. **B)** Distance to the bottom of the assay in each 30 seconds for the three genotypes. **C)** Fraction of time in the bottom third and **D)** top halve visits for the three genotypes during the novel tank diving assay. Graphs show mean ± SEM

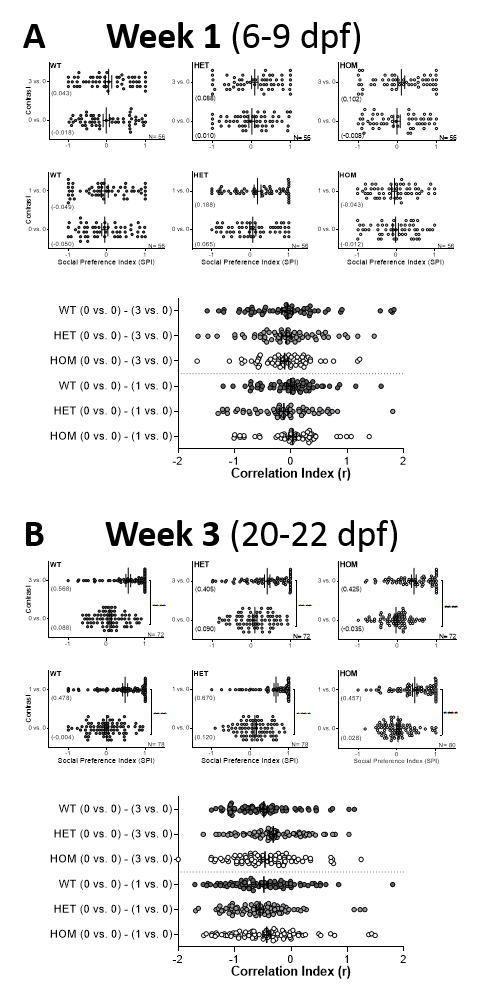

Supplementary figure 6. **Ontogeny of sociability in zebrafish *baz1b^del44^* LoF at week 1 and week 3**. For each week, graphs show change in social preference index (SPI) between basal conditions (0 vs. 0) and contrast (0 vs. 3 or 0 vs. 1) for either WT, HET or HOM. N is included in each graph. Right panels show the comparison of the correlation indexes for all the contrast at the corresponding week. A) week 1 (from 6 to 9 dpf), B) week 3 (from 20 to 23 dpf). Graphs show individual values and mean (also between brackets for SPI) ± SEM. In all cases: **** *p* < 0.001

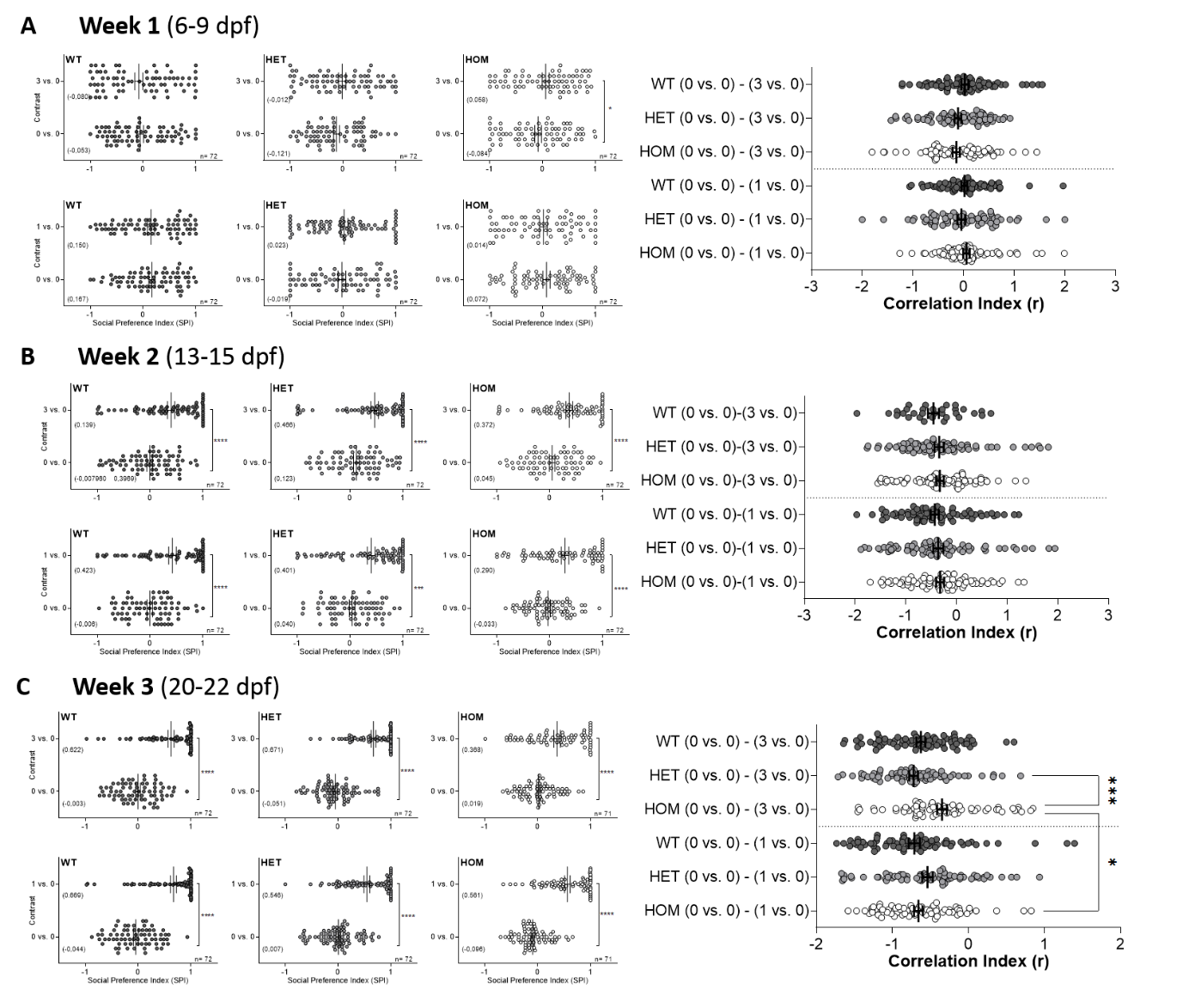

Supplementary figure 7. **Loss of *baz1b^ins35^* disturbs the ontogeny of sociability in zebrafish**. For each week, graphs show change in social preference index (SPI) between basal conditions (0 vs. 0) and contrast (0 vs. 3 or 0 vs. 1) for either WT, HET or HOM. N is included in each graph. Right panels show the comparison of the correlation indexes for all the contrast at the corresponding week. A) week 1 (from 6 to 9 dpf), B) week 2 (from 13 to 15 dpf), C) week 3 (from 20 to 22 dpf). Graphs show individual values and mean (also between brackets for SPI) ± SEM. In all cases: * *p* < 0.05; ** *p* < 0.01; **** *p* < 0.001
